# Overcoming systematic data biases enables accurate prediction of enzyme *k*_cat_ fold-changes for computational protein design

**DOI:** 10.64898/2026.01.23.701068

**Authors:** Yvan Rousset, Alexander Kroll, Martin J. Lercher

## Abstract

Machine learning is increasingly used to guide protein engineering by predicting how mutations affect desired properties. Recent models for the turnover number (*k*_cat_) of enzymes report high accuracy, suggesting that mutation effects can be inferred directly from protein sequence. However, these approaches are typically evaluated on heterogeneous datasets of enzyme variants, where closely related sequences and systematic reporting patterns may confound model performance. A central challenge is therefore to determine whether current models truly capture mutation-specific effects or instead exploit statistical regularities in the data. Here we show that much of the reported accuracy in mutant *k*_cat_ prediction arises from two pervasive biases: variants of the same enzyme occupy a narrow activity range, and mutations within a group often share a common direction of change. Simple baselines that exploit these biases match or exceed the performance of existing models, indicating that high apparent accuracy does not imply mechanistic understanding. To address this limitation, we introduce a bias-aware framework that reformulates prediction as a pairwise fold-change task and evaluates performance on unseen mutant–mutant pairs, thereby isolating mutation-specific signal. A proof-of-principle implementation explains approximately one-third of the variance under these conditions and outperforms existing models on leakage-controlled benchmarks. More broadly, this work establishes a general strategy for evaluating and modeling mutation effects in biochemical datasets, with implications for protein engineering and related fields.

## Introduction

Designing proteins with tailored functions is a central goal of biotechnology^1–6^. For enzymes, a key target is the turnover number, *k*_cat_, which quantifies the maximal number of substrate molecules converted per active site per unit time and is therefore directly relevant to biocatalyst optimization, enzyme engineering, and metabolic modeling. Because experimental measurements of *k*_cat_ are laborious, sparse, and heterogeneous, there is growing interest in using machine learning to predict kinetic parameters from sequence and biochemical context.

A major development enabling these approaches has been the emergence of protein language models, which learn general-purpose representations of protein sequences from large sequence databases^7–11^. These representations have proved useful across broad protein prediction tasks and can also be used for more specialized enzyme-function prediction. For *k*_cat_ prediction, however, relying on the enzyme sequence alone is likely to be suboptimal, because turnover depends on the substrate and, more generally, on the biochemical reaction being catalyzed. Several *k*_cat_ predictors combine protein representations with substrate representations to predict enzyme–substrate-specific turnover numbers^12–15^, whereas other approaches incorporate more complete reaction descriptions^16^. The broader problem of jointly representing proteins and small molecules has also been reviewed in detail^17^.

Although these models report apparently strong performance and have been proposed as tools for high-throughput kinetic annotation and enzyme engineering, current approaches might fail to capture mutation-specific effects^18^. For enzyme engineering, this distinction is critical. The practically relevant question is often not the absolute *k*_cat_ value of an isolated enzyme–substrate pair, but how specific mutations change activity relative to a known parent enzyme. Distinguishing this mutation-specific signal from broader enzyme-, substrate-, or reaction-level differences is therefore essential for evaluating whether models can guide enzyme optimization.

We show that the failures of existing predictors for mutant *k*_cat_ arise from systematic biases in mutant kinetic datasets. First, *k*_cat_ values for variants of the same enzyme cluster within a narrow range, so that correctly identifying their enzyme identity yields deceptively high accuracy (range bias). Second, reported mutations within a given enzyme are often consistently activating or deactivating, allowing models to succeed by predicting the prevailing direction of change (directional bias). Together, these effects inflate performance metrics and create the illusion that models capture mutation-specific effects far more than they actually do.

To eliminate these shortcuts, we reformulate *k*_cat_ prediction as a pairwise task, predicting fold changes between variants rather than absolute values and validating on unseen mutant–mutant pairs. This removes the incentive for models to exploit group-level biases, forcing them to focus on mutation-induced differences instead. As a proof of principle, we implement this formulation in a simple prediction model, FCKcat, and evaluate it on leakage-controlled benchmarks using only previously unseen mutants. Under these conditions, apparent performance drops substantially, revealing the true difficulty of the problem, while FCKcat captures a substantial fraction of mutation-specific variation and outperforms existing methods.

## Results

To investigate mutant kinetic behavior, we assembled a curated dataset of enzyme variants from the BRENDA^19^ and SABIO-RK^20^ databases (Methods), comprising 8 289 measurements across 7 011 unique sequences. We grouped sequences by enzyme–reaction pair (shared UniProt ID and reaction), yielding 1 526 groups (2–81 sequences each). This grouping separates enzyme-level variation from mutation-specific effects and forms the basis for all analyses. From these groups, we generated sequence pairs for model training and evaluation, using a partitioning strategy that ensures no mutant sequence appears in both training and test sets, thereby preventing direct memorization of variants.

### Mutant *k*_cat_ data exhibits two distinct biases

If all variants of an enzyme have similar catalytic rates, a model can appear accurate by simply predicting a typical value for that enzyme. We first quantified the extent to which this effect occurs. Across the full dataset, *k*_cat_ values span almost 14 orders of magnitude (∼ 10^−7^ to 10^7^ s^−1^; Fig. S1a). However, within individual enzyme–reaction groups, this diversity largely disappears: most variants differ by less than a factor of ∼5 (mean within-group standard deviation on log_10_-scale: 0.69; Fig. S1b). Consistent with this observation, enzyme–reaction group identity explains 66.1% of the variance in log-transformed *k*_cat_ values (intraclass correlation coefficient *ICC* = 0.661; Fig. 1a), indicating that most variation reflects differences between enzymes rather than differences between mutations within the same enzyme.

**Figure 1.**
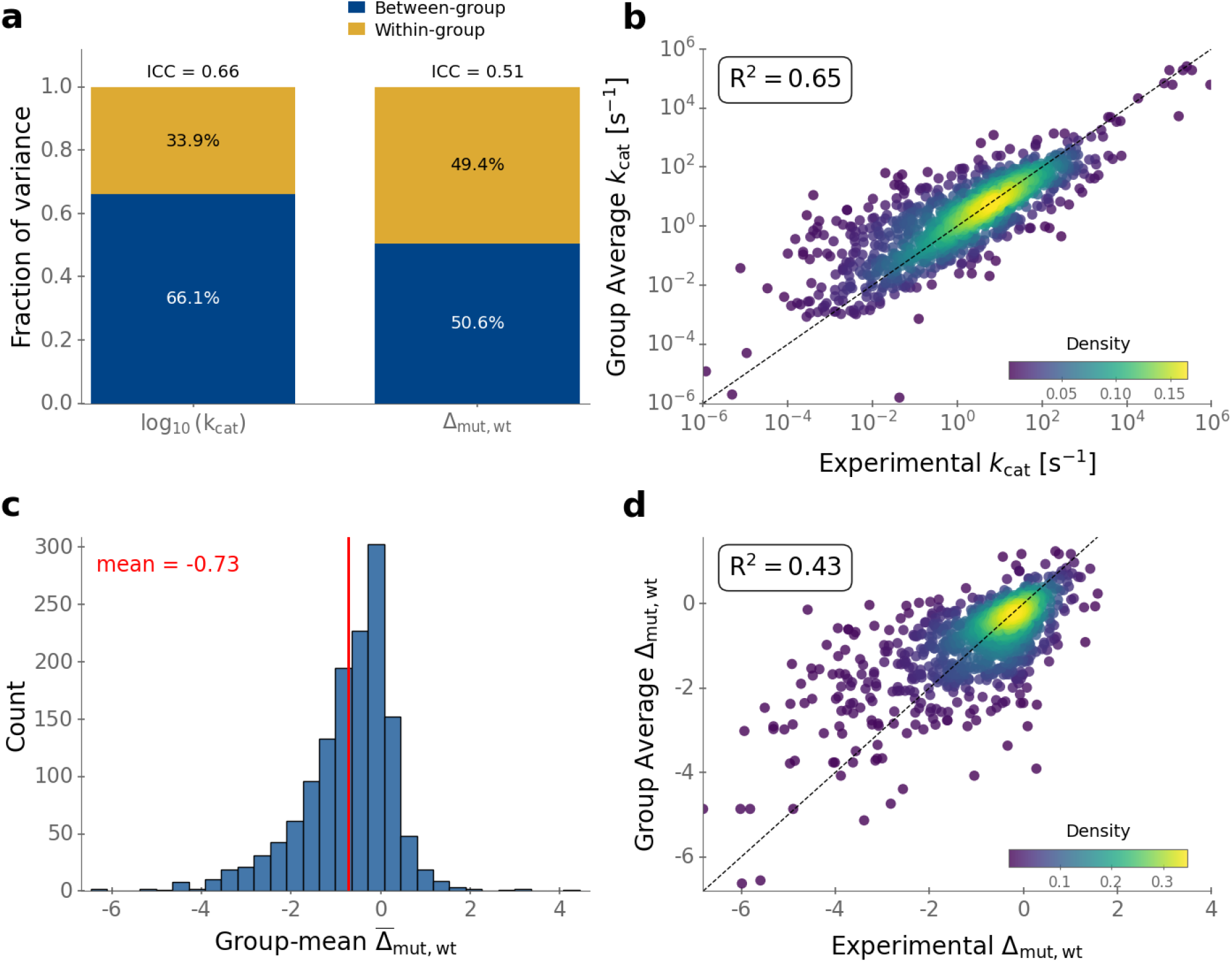
Apparent predictive accuracy arises from enzyme–reaction group-level biases, not mutation-specific information. **a**, Variance partitioning using random-intercept mixed-effects models. Stacked bars show the fraction of variance attributable to between-group and within-group sources. Enzyme–reaction group identity accounts for 66% of the variance in absolute log_10_ *k*_cat_ (left; *range bias*) and 50% in mutant–wild-type fold changes (right; *directional bias*), indicating that much of the apparent predictability arises from group-level structure rather than mutation-specific effects. **b**, Baseline for absolute *k*_cat_ prediction exploiting *range bias*. Each test mutant is assigned the mean log_10_ *k*_cat_ of all training variants (wild-type and mutant) from the same enzyme–reaction group. Despite ignoring mutation-specific information, this baseline achieves high accuracy. **c**, Distribution of directional bias across enzyme–reaction groups, defined as the mean log_10_ ratio between mutant and wild-type *k*_cat_. Positive values indicate that mutations predominantly increase *k*_cat_, negative values indicate decreases. Approximately 20% of groups show a consistent increase, illustrating strong group-level skew in the direction of mutational effects. **d**, Baseline for *k*_cat_ fold-change prediction exploiting *directional bias*. Each mutant–wild-type pair is assigned the mean log-fold change Δ_mut,wt_ from training mutants in the same group. This baseline accurately predicts both magnitude and direction of change without using mutation-specific information, demonstrating that high predictive accuracy can be achieved without modeling mutation-specific effects. Density coloring in (b,d) indicates data point concentrations, dashed diagonals indicate perfect predictions.

To assess how easily a model can exploit this *range bias*, we considered a simple group-mean baseline. For each test mutant, we predicted its log_10_ *k*_cat_ as the mean value of all training variants from the same enzyme–reaction group, without using any information about the specific amino acid changes. This trivial predictor achieves *R*^2^ = 0.65 (Fig. 1b), matching or exceeding the performance reported for state-of-the-art models such as EITLEM, UniKP, DLKcat, and TCNeKP^12–15^. Restricting the mean to other mutants within the group further increases performance to *R*^2^ = 0.72. Thus, the reported accuracy of current models can be reproduced by baselines that simply exploit the narrow range of *k*_cat_ values within enzyme groups, without learning mutation-specific effects. This group-mean predictor therefore defines a natural null model: any method that does not outperform it is unlikely to meaningfully capture mutation-specific effects.

Beyond this clustering of *k*_cat_ values, mutant datasets exhibit a second, equally problematic bias related to the direction of mutational effects. If mutations were sampled without bias, increases and decreases in *k*_cat_ would occur with equal frequency. Instead, we observe that mutations within a given enzyme–reaction group are often strongly skewed, with most variants either consistently increasing or consistently decreasing activity (Fig. 1c). This *directional bias* likely reflects experimental practice and reporting, where studies typically focus on identifying either activating or inactivating mutations.

To isolate this effect, we consider fold changes between variants. For two sequences (*a, b*) from the same enzyme–reaction group, we define the log-fold change as

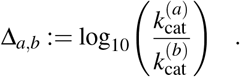

When expressing mutant effects as log-fold changes relative to the wild-type, enzyme–reaction group identity still explained 50.6% of the variance (ICC = 0.506; Fig. 1a), indicating that directional trends are strongly group-specific.

We next tested whether this bias alone is sufficient to yield high predictive accuracy. For each enzyme–reaction group with wild-type wt, we computed the mean fold change Δ_mut,wt_ across training mutants and used this single group-level value to predict the fold change for each test mutant. This baseline achieves *R*^2^ = 0.43 (Fig. 1d) and correctly predicts the direction of change in 83.6% of cases, again without using any mutation-specific information. This directional baseline defines a second null model: accurate prediction of the direction or magnitude of mutational effects does not imply mechanistic understanding unless it exceeds this group-level expectation.

Together, these results show that high apparent accuracy in predicting mutant effects can arise entirely from dataset biases. Models can achieve strong performance by exploiting the range and directionality of *k*_cat_ values within enzyme groups, without capturing the biochemical consequences of individual mutations. The corresponding group-level baselines therefore define natural null models for mutant *k*_cat_ prediction: any method that fails to outperform them does not capture mutation-specific effects. Consequently, standard performance metrics on absolute *k*_cat_ prediction are insufficient to assess whether a model has learned the biochemical consequences of mutations. This insight motivates a reformulation of the prediction task that explicitly removes these shortcuts.

### A pairwise fold-change model circumvents systematic biases

Because absolute *k*_cat_ prediction can be solved by exploiting group-level biases, it does not require learning mutation-specific effects. To eliminate these shortcuts, we reformulate the task as predicting the log-fold change Δ_*a,b*_ between two sequences (*a, b*) from the same enzyme–reaction group. This pairwise formulation offers three advantages:

- **Signal isolation:** the shared group-level scale cancels out, forcing the model to focus on mutation-induced differences rather than typical enzyme activity.
- **Data expansion:** for a group of size *n*, the number of training examples increases from *n* to *n*(*n* −1)/2; for our dataset, this is an expansion from 8 289 entries to 28 809 pairs.
- **Predictive flexibility:** pairwise predictions can be combined across multiple reference sequences to obtain more robust estimates for new variants.

As a proof of principle, we implemented this formulation in a simple model, FCKcat. Enzyme sequences were encoded using ESM-2^9^. For each sequence pair (*a, b*), we concatenated ESM-2 representations from each sequence:

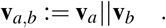

A gradient-boosting model was trained to predict the corresponding log-fold change Δ_*a,b*_ (Fig. 2a).

**Figure 2.**
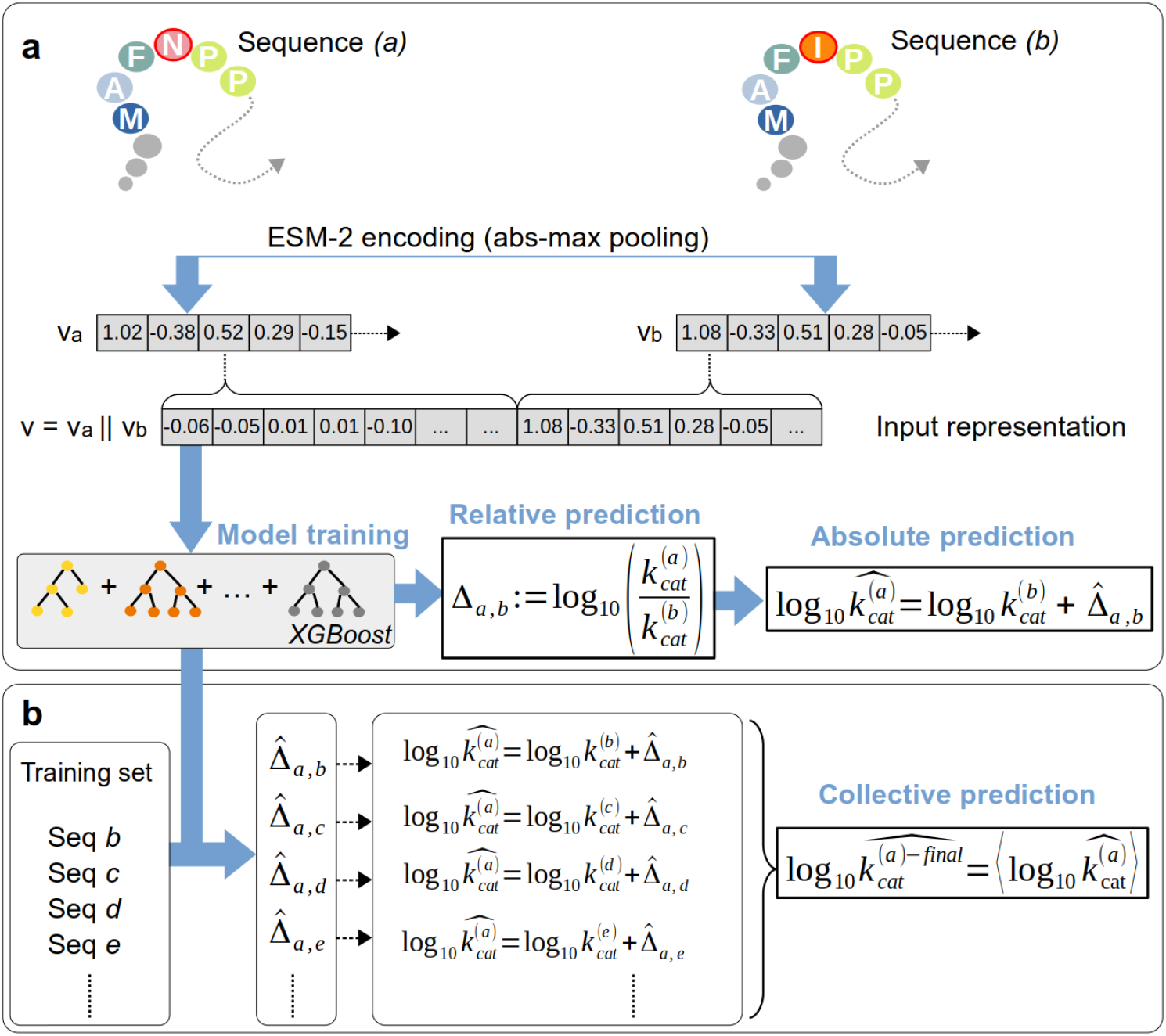
Pairwise model for predicting relative *k*_cat_ changes between enzyme variants. **a**, Enzyme sequences (a) and (b) are encoded into a vector representation **v**_(*a*)_ and **v**_(*b*)_ which account for the mutation-induced difference. A gradient boosting model predicts the corresponding log-fold change Δ_*a,b*_. **b**, For a given enzyme–reaction group, predictions for a new variant can be obtained by pairing it with multiple reference sequences from the same group and averaging the resulting estimates to obtain a more accurate collective prediction.

Standard evaluation protocols, in which wild-type anchors are present in both training and test sets, allow models to exploit group-level biases. To directly assess mutation-specific prediction, we introduce a stringent evaluation protocol that removes all sources of such shortcut learning (Methods). Wild-type sequences were excluded from validation and test sets, forcing the model to predict functional differences between pairs of entirely unseen mutants. Under these bias-controlled conditions, trivial baselines collapse to zero performance.

In this setting, FCKcat explains approximately one-third of the variance (*R*^2^ = 0.31 in cross-validation and *R*^2^ = 0.31 on an independent test set; Fig. 3a). Although lower than values reported for absolute *k*_cat_ prediction, this performance corresponds to a substantially harder task in which shortcut learning is no longer possible. The remaining signal therefore reflects mutation-specific variation beyond group-level effects and provides a lower bound on the mutation-specific information contained in current datasets. When mutant–wild-type pairs were included, performance was *R*^2^ = 0.35 on the full test set and *R*^2^ = 0.27 on mutant–wild-type pairs only (Fig. S2). The latter value should be interpreted with caution because all mutant–wild-type pairs were ordered with the mutant as the first sequence, producing a non-centered target distribution that can deflate *R*^2^.

**Figure 3.**
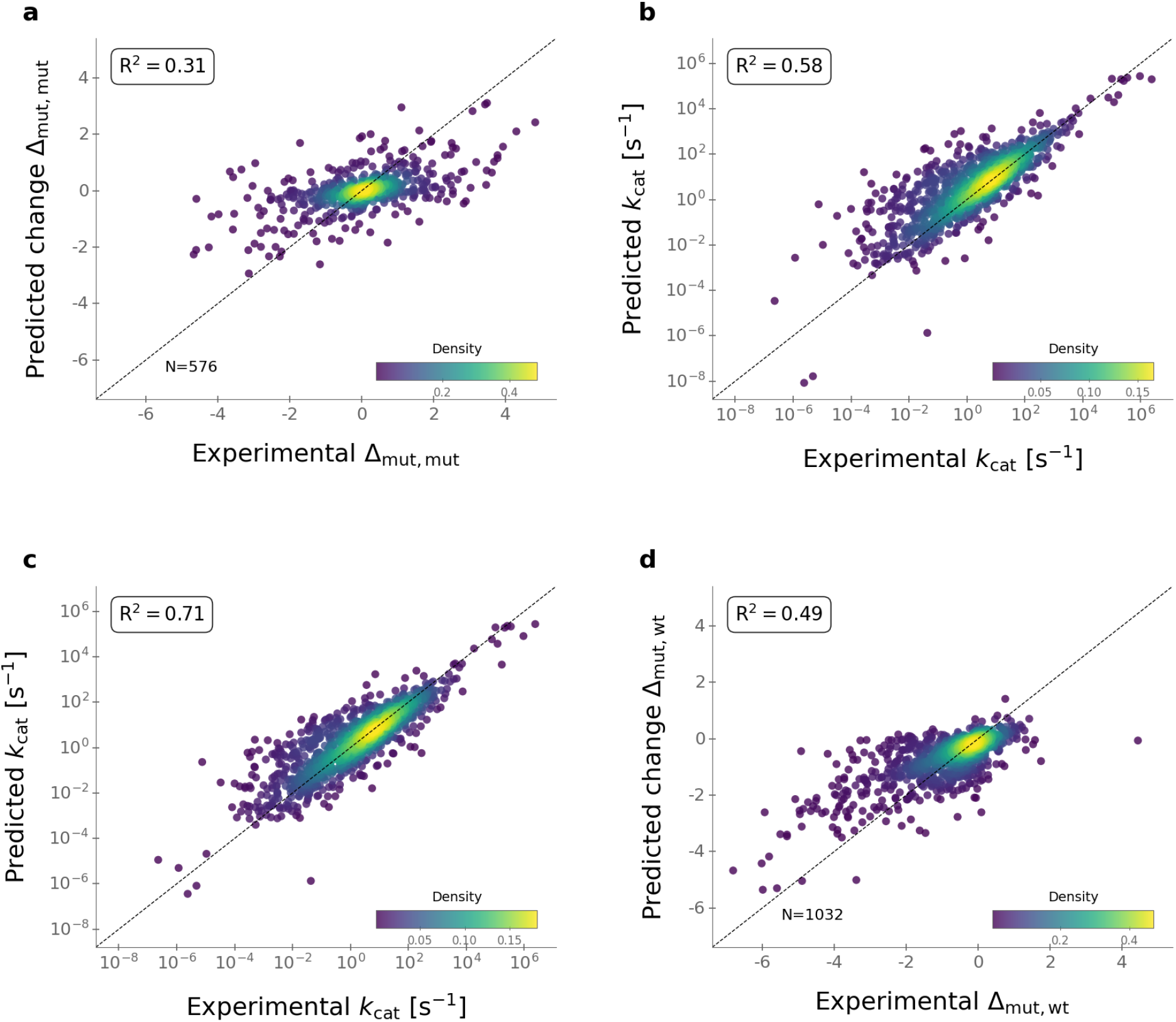
From pairwise fold-change prediction to direct and collective reconstruction of absolute *k*_cat_. **a**, Prediction of log-fold changes Δbetween pairs of unseen mutants. This setting removes group-level shortcuts and reflects mutation-specific predictive performance. **b**, Direct reconstruction of absolute *k*_cat_ values by anchoring predicted fold changes to a measured reference. This reintroduces group-level information and inflates apparent accuracy. *R*^2^ values in **b, c** are calculated on log-transformed data. **c**, Collective reconstruction of absolute *k*_cat_ values obtained by averaging predictions across multiple reference sequences from the same enzyme–reaction group, reducing variance and improving accuracy in practice. **d**, Recovery of mutant–wild-type fold changes from collectively reconstructed absolute values. Density coloring in all panels indicates data point concentrations, dashed diagonals indicate perfect predictions.

### Collective predictions boost prediction accuracy

While our model is trained and evaluated on fold changes, thereby isolating mutation-specific effects, absolute *k*_cat_ values can be reconstructed by anchoring a predicted fold change to a known reference:

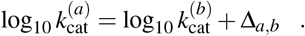

This reconstruction reintroduces group-level information through the reference value and therefore partially restores range and directional biases, leading to inflated performance metrics. On our test set of wild-type–mutant pairs, this simple anchoring yields *R*^2^ = 0.58 (Fig. 3b). The reduced performance compared to the group-level baseline (Fig. 1b) likely reflects noise in wild-type *k*_cat_ measurements.

To reduce reliance on any single reference, we developed a collective inference strategy that exploits the model’s pairwise structure. For a given mutant, we generate multiple independent estimates by pairing it with all available reference sequences (wild-type or mutant) within the same enzyme–reaction group. Averaging these predictions (Fig. 2b) reduces variance and yields *R*^2^ = 0.71 for absolute *k*_cat_ predictions and *R*^2^ = 0.49 for fold changes relative to the wild-type (Fig. 3c,d).

Importantly, this improvement reflects variance reduction rather than additional mutation-specific information: by aggregating multiple pairwise estimates, the model leverages shared group context to average out noise in reference *k*_cat_ measurements and prediction noise. Accordingly, collective prediction should be viewed as a practical strategy for improving absolute *k*_cat_ estimates in data-rich settings, rather than as a bias-free measure of mutation-specific predictive performance.

Formally, this procedure can be viewed as a *U* -statistic: the final prediction is obtained by applying a pairwise function to sequence pairs and averaging the resulting values^21^. Notably, the model retains a non-negligible predictive performance of approximately 20% even when no reference variant is available, and performance increases progressively with the number of characterized reference variants. In practice, even a small number of reference variants is sufficient to improve predictions, with as few as two characterized mutants substantially increasing accuracy (Fig. S3).

### FCKcat predicts the direction of *k*_cat_ change with high confidence for large effects

Accurately predicting whether a mutation increases or decreases catalytic activity is a key requirement for practical enzyme design. FCKcat achieves a sign accuracy of 69% for directional change between two unseen mutant sequences where a random guess would achieve 50% (the mutant–mutant test pairs are sign-balanced by construction, so 50% is the chance rate). We therefore examined how directional reliability depends on the predicted effect magnitude 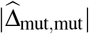. While overall sign accuracy is 69%, it increases sharply with predicted magnitude, reaching 90% for mutations predicted to change activity by more than one order of magnitude (Fig. S4), and 100% for two orders of magnitude. This trend indicates that large predicted effects are highly reliable, whereas small predicted changes are more susceptible to noise and residual bias.

### FCKcat outperforms existing models on leakage-controlled, bias-aware benchmarks

Recent models for absolute *k*_cat_ prediction report high performance, but these evaluations are susceptible to the biases identified above and to training–test overlap. To enable a fair comparison, we performed leakage-controlled evaluations against DLKcat^14^ and EITLEM^12^ (Methods; UniKP^13^ was excluded due to the unavailability of its original splits). For each comparison, we constructed independent test sets consisting exclusively of mutant–mutant pairs in which neither sequence appeared in the corresponding training datasets. Because these leakage-controlled comparisons rely on slightly different test sets, the FCKcat values reported here differ slightly from those obtained on the full FCKcat test set.

Under these bias-aware conditions, FCKcat consistently outperformed DLKcat and EITLEM across all metrics (Fig. 4a,b,c), including *R*^2^, Pearson correlation, directional accuracy, and Matthews correlation coefficient (MCC). The performance gap was most pronounced for MCC, indicating that FCKcat provides more balanced and reliable predictions of the direction of *k*_cat_ change.

**Figure 4.**
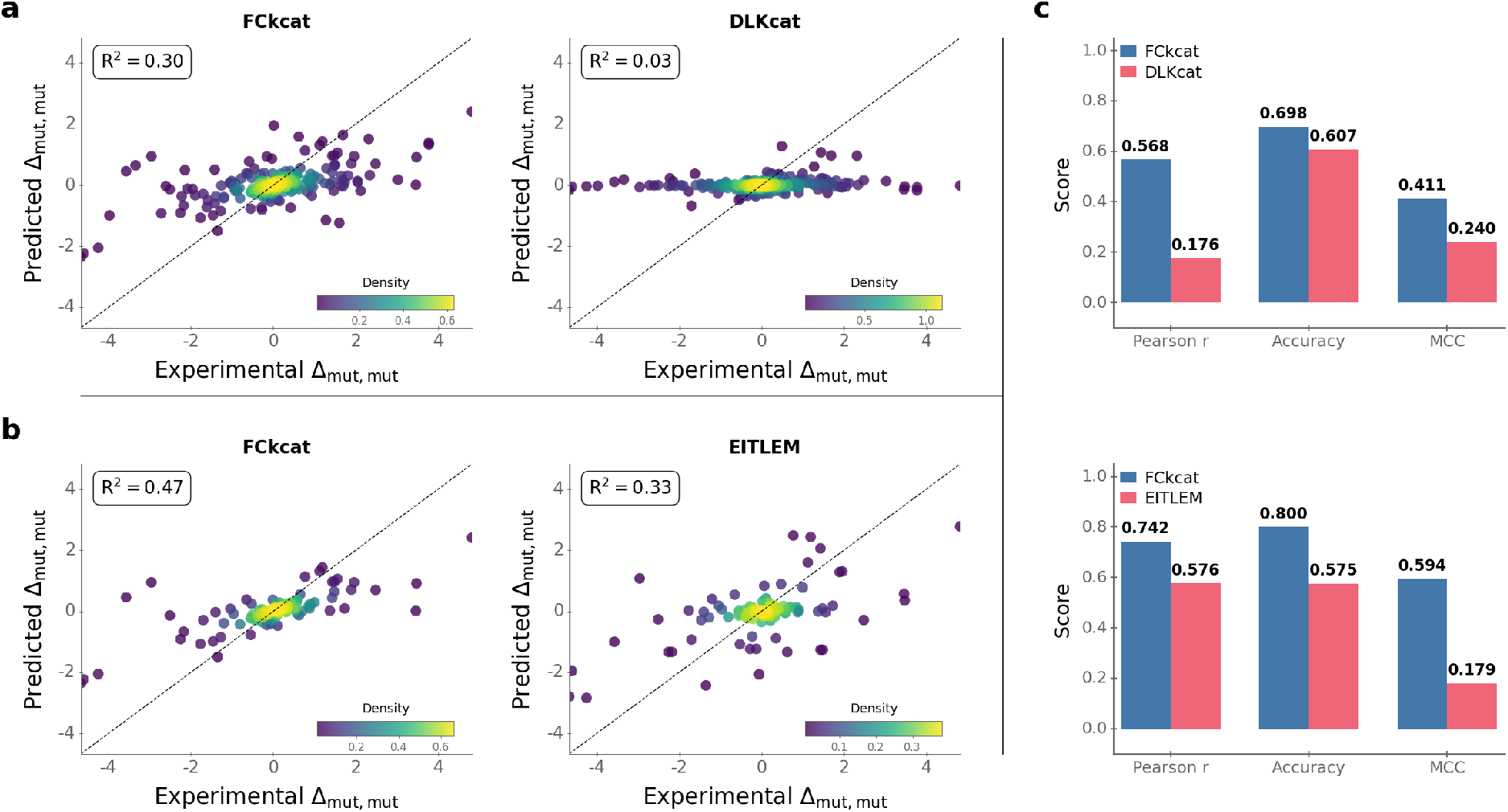
FCKcat outperforms existing models on mutation-specific prediction under leakage-controlled, bias-aware evaluation. **a**,**b**, Predicted versus experimental log-fold changes Δ_*a,b*_ for unseen mutant–mutant pairs, evaluated on the leakage-controlled test set intersections between FCKcat and DLKcat (top row) and EITLEM (bottom row). Density coloring indicates point density; dashed diagonals indicate perfect agreement. **c**, Quantitative performance metrics for the same comparisons, including Pearson correlation (*r*), directional accuracy, and Matthews correlation coefficient (MCC). FCKcat consistently outperforms both models across all metrics.

These results demonstrate that, when evaluated under conditions that eliminate shortcut learning, FCKcat more effectively captures mutation-specific effects than existing models, whose performance is strongly influenced by group-level biases and data leakage.

## Discussion

In this study, we identify two systematic biases in mutant *k*_cat_ datasets that can give rise to misleadingly high predictive performance. First, *k*_cat_ values for variants of a given enzyme typically cluster within a narrow range. As a result, predicting values close to the wild-type yields deceptively strong accuracy, as noted previously^18^. We show that a simple group-mean baseline exploiting this range bias matches the performance of existing models despite ignoring mutation-specific effects. Second, mutations within an enzyme–reaction group are often directionally skewed, predominantly increasing or decreasing activity. A corresponding baseline that predicts the typical direction of change can therefore achieve high apparent accuracy without capturing mutation-specific information. Together, these effects demonstrate that performance on absolute *k*_cat_ prediction largely reflects group-level structure rather than the biochemical consequences of individual mutations.

These biases are not specific to our dataset but are also present in datasets used by other models^12,14^ (Fig. S5). They arise naturally from how enzyme kinetics data are generated and reported. Variants of a given enzyme share the same catalytic mechanism and structural scaffold, constraining *k*_cat_ to a narrow functional range and giving rise to range bias. In addition, mutant datasets are typically generated through targeted studies aimed at enhancing or disrupting activity, leading to systematic directional skew. Consequently, most enzyme-variant datasets will exhibit strong group-level structure, enabling models to achieve high apparent accuracy without learning mutation-specific effects.

We propose to eliminate these shortcuts by reformulating the task as the prediction of fold changes between pairs of sequences. This removes the shared group-level scale and forces the model to encode mutation-specific effects. Combined with a leakage-controlled evaluation on unseen mutant–mutant pairs, this approach prevents reliance on group-level directionality. Under these conditions, our proof of principle model FCKcat explains approximately one-third of the variance in mutant–mutant fold changes, outperforming previous methods and providing a conservative lower bound on the mutation-specific signal contained in current datasets.

For practical applications requiring absolute *k*_cat_ estimates, the pairwise framework can be extended to collective inference, formally a *U* -statistic. Our results show that aggregating pairwise estimates reduces variance and improves accuracy, yielding *R*^2^ = 0.71 with our simple model. Importantly, this improvement reflects variance reduction and reintroduction of group-level information, rather than increased mutation-specific signal.

Under leakage-controlled and bias-aware evaluation, the pairwise framework enables meaningful comparison of models. Within this setting, FCKcat outperforms the much more complex models DLKcat and EITLEM on fold-change prediction, indicating that it captures mutation-specific effects more effectively. More broadly, these results highlight that improvements in model architecture are only informative when evaluated within a bias-aware framework that eliminates shortcut learning.

Kinetic data from BRENDA and SABIO-RK contain substantial noise, including erroneous data transfer from the literature (affecting ∼ 20% of entries^22,23^), misannotated mutations, and inconsistencies in mutation formatting. In addition, experimental conditions such as temperature, pH, and cofactors vary widely and are not fully captured by current feature sets, further limiting achievable accuracy. Fold-change prediction partially mitigates these issues: measurements within a group are often obtained under comparable conditions, allowing systematic errors to cancel when taking ratios. Nevertheless, our estimate of *R*^2^ *≈* 0.30 likely underestimates the intrinsic predictability of mutational effects in cleaner datasets.

A key application of our framework is the identification of mutations that enhance catalytic activity. Although trained predominantly on non-enhancing variants, FCKcat reliably identifies high-confidence improvements: the probability that a predicted increase is correct rises sharply with predicted effect size. This enables practitioners to trade coverage for reliability by thresholding predictions. In practice, the model can be used to prioritize candidate mutations or small combinatorial libraries for experimental validation, reducing search space and accelerating enzyme optimization^24^. As additional mutants are experimentally characterized, prediction accuracy further improves through collective inference.

Our architecture is intentionally simple, relying on gradient boosting over concatenated absolute max pooling representations from pre-trained protein sequence embeddings. Interestingly, an element-wise difference representation performed nearly as well despite omitting the individual enzyme embeddings, suggesting that pairwise differences in ESM representation space already retain much of the sequence-derived information relevant for fold-change prediction. Further improvements could be achieved by incorporating the reference sequence and reaction context more explicitly, adding physicochemical features, accounting for experimental conditions, and fine-tuning representations for fold-change prediction. We leave these directions for future work.

Taken together, our results define a general framework for mutation-effect prediction: models should be trained on relative, pairwise tasks and evaluated under leakage-controlled conditions that remove group-level shortcuts. Within this framework, performance reflects mutation-specific signal rather than dataset structure. While we here focus on *k*_cat_, the same approach is likely to improve prediction of other properties such as binding affinities, Michaelis constants (*K*_M_), catalytic efficiency (*k*_cat_*/K*_M_), and stability.

## Methods

### Software and Code Availability

All analyses were performed in Python (v3.9). We fitted the gradient boosting models using the library XGBoost. The code and data used to generate the results of this paper, in the form of Jupyter notebooks, is available from https://github.com/yvanrousset/FCKcat.

### Variance partitioning with mixed-effects models

To quantify how much variation in catalytic rates is attributable to enzyme–reaction group structure, we fitted random-intercept linear mixed-effects models on the original, non-paired measurements. For the analysis of absolute catalytic rates, the response variable was the log-transformed geometric mean catalytic rate, *y*_*ig*_ = log_10_(*k*_cat,*ig*_), for observation *i* in enzyme–reaction group *g*, modeled as

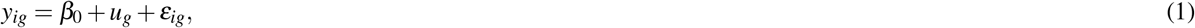

where *β*_0_ is the global intercept, *u*_*g*_ is the random intercept for group *g*, and *ε*_*ig*_ is the residual error term.

To quantify directional bias, we constructed one mutant–wild-type log-fold change per mutant,

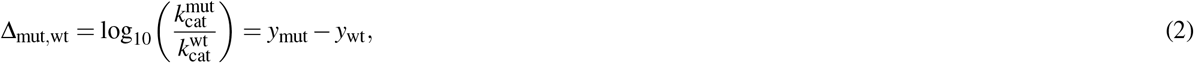

using the wild-type measurement from the same enzyme–reaction group as reference. Wild-type rows themselves were excluded from this analysis, and groups lacking a valid wild-type reference were omitted. We then fitted an analogous random-intercept model,

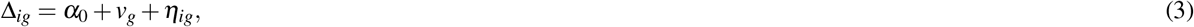

where Δ_*ig*_ is the mutant effect for mutant *i* in group *g, v*_*g*_ is the group-level random intercept, and *η*_*ig*_ is the residual error.

Variance partitioning was summarized using the intraclass correlation coefficient (ICC),

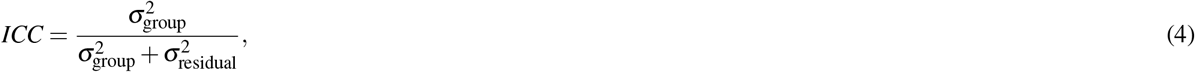

which gives the proportion of total variance attributable to enzyme–reaction group identity. Models were fitted in Python using statsmodels with restricted maximum likelihood (REML). Sensitivity analyses excluding singleton groups gave nearly identical estimates.

### Data extraction and preprocessing

#### Source extraction and preliminary normalization

Raw kinetic measurements were obtained from BRENDA^19^ and SABIO-RK^20^. Because the two resources differ in data format and annotation structure, entries were first extracted separately and standardized before cross-database harmonization.

For BRENDA, we processed the nested JSON export, in which EC numbers are used as top-level keys, into a tabular format and retained EC number, organism, UniProt identifier^25^, substrate/product annotations, free-text commentary, and *k*_cat_ values. Variant status was inferred from the commentary field using keywords such as “wild-type”, “mutant”, “mut”, and “recombinant”, as well as mutation patterns such as “A182Y”. Entries lacking explicit variant keywords were provisionally treated as wild-type and subsequently checked during mutation curation. After removal of missing values, the BRENDA dataset comprised 21,032 provisional wild-type and 13,831 provisional variant entries.

For SABIO-RK, we retrieved *k*_cat_ entries together with organism, UniProt identifier, EC number, substrate/product annotations and, when available, KEGG reaction identifiers. Mutation information was extracted from the “enzyme variant” field and curated as described below. Entries lacking UniProt identifiers were excluded, units were homogenized to s^−1^, and reported zero values were reassigned to 10^−6^ s^−1^ to enable log_10_ transformation.

#### Mutation parsing and sequence reconstruction

For both BRENDA and SABIO-RK, variant annotations, including mutation information, are recorded in free-text fields. Because reporting formats vary widely, including heterogeneous notation styles, mutation types, inconsistent vocabularies and occasional errors, we adopted a semi-automated curation workflow. We first applied pattern-matching rules to extract common mutation notations, then manually reviewed the remaining ambiguous entries, approximately 1,500, and, when necessary, consulted the original publications (70). Manual curation focused on two recurrent failure modes: (i) wild-type enzymes to which spurious mutations had been assigned, typically due to parsing experimental conditions or molecule names, and (ii) mutant entries for which no mutation was captured automatically, most commonly because the change involved deletions or insertions with highly variable notation. We also removed strings that were falsely identified as mutations but clearly corresponded to other entities, such as “H2O”, “H4C” and “P2A”, or recurrent short patterns, such as “L1A” and “K3F”, that often denote variant names rather than true residue changes.

To validate and reconstruct sequences, we downloaded each wild-type sequence from UniProt and applied the curated mutations in silico. Substitution strings were parsed into original residue, if provided, position and mutant residue, sorted, and applied sequentially. When the original residue was specified, we verified it against the wild-type sequence; mismatches were flagged. For single mismatches, we attempted correction by checking adjacent positions; for multiple mismatches, we inferred a consistent positional offset when supported by the surrounding context. We applied the same procedure for single-residue deletions and insertions. Entries with larger indels or insufficient positional detail were retained only when the resulting sequence could be reconstructed unambiguously; otherwise, they were excluded.

In total, 1,300 entries were excluded because their curated mutation annotations could not be reconciled with the associated UniProt wild-type sequence.

#### Harmonization, reaction mapping and replicate aggregation

Substrates and products were mapped to KEGG compound identifiers and, when direct mapping was unavailable, to PubChem compound identifiers and back to KEGG using *mbrole2*^26^. KEGG reaction identifiers were then assigned by matching EC numbers and substrate sets against the KEGG database, retaining only entries that could be mapped to curated (“natural”) KEGG reactions.

After reaction mapping, BRENDA and SABIO-RK entries were concatenated and cross-source duplicates were removed, defined as entries with the same UniProt ID, curated mutation annotation, substrate/reaction assignment and *k*_cat_ value (690 entries removed). Entries were then grouped by UniProt ID, KEGG reaction ID and curated mutation annotation. When multiple *k*_cat_ values remained for the same group, they were summarized by their geometric mean. Enzymes associated with multiple UniProt identifiers were excluded to avoid ambiguous assignment in multi-subunit complexes.

Reaction representations were encoded as reaction SMILES by mapping each KEGG metabolite identifier to ChEBI/PubChem via the KEGG and ChEBI APIs, using *bioservices*^27^, and retrieving SMILES or InChI strings; InChI strings were converted to SMILES using RDKit. Reactions containing undefined compounds or lacking valid molecular structures for any substrate or product were removed. Finally, we retained only enzyme–reaction groups containing at least one wild-type and one mutant measurement, enabling supervised learning on paired targets 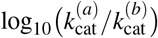.

After all extraction, curation, harmonization and filtering steps, the final dataset contained 8,289 entries, including 1,391 wild-type and 6,898 mutant measurements. The mutant measurements comprised 6,787 entries with one or more substitutions, 103 with one or more deletions, 4 with one or more insertions, 2 with one or more additions, and 2 with both deletions and substitutions.

### Datasets and splitting strategies

The final dataset comprised 8,289 *k*_cat_ values, covering 7,011 unique sequences (1,085 unique wild-type and 5,926 unique mutant), associated with 956 unique reactions. Entries were grouped by unique enzyme–reaction pair (a wild-type and its mutants), yielding 1,526 groups (sizes 2–81). We define an enzyme–reaction group as all entries sharing the same UniProt ID and the same KEGG reaction ID (wild-type plus all associated mutants). This grouping prevents comparing variants across distinct reactions and ensures fold changes are defined only within a fixed enzyme–reaction context. Our splitting strategy was guided by the most relevant application scenario: because mutants are defined relative to a wild-type and an important use case is to identify mutants with enhanced kinetics, we aim to predict whether a new mutant has a higher *k*_cat_ than its wild-type, and by how much.

To increase dataset size while preserving realistic evaluation, mutants within each group were randomly distributed across six splits: five cross-validation (CV) folds and one external test set, as evenly as possible. The wild-type was included in every split that contained at least one mutant (see Methods). For example, in a group with one wild-type and six mutants, one mutant was assigned to each CV fold and one to the test set; each split thus contained the same wild-type and exactly one mutant. During training, four folds were pooled and all unordered sequence pairs within those folds were formed, yielding *n*(*n* − 1)/2 pairs for *n* sequences in the pooled training split; in the six-mutant example this gives 5 · 4/2 = 10 training pairs, with one pair left for validation in the held-out fold and one pair in the external test set. Across the five CV folds, this procedure produced a total of 15 training pairs and 1 test pair for that example. We evaluated two variants of this scheme: (i) keeping wild-type–mutant pairs in validation/test, and (ii) removing all wild-type–mutant pairs from validation/test to reduce bias and focus on mutant–mutant comparisons (Fig. 5); the latter is more challenging because many groups have few or no mutant–mutant pairs assignable to validation or test.

**Figure 5.**
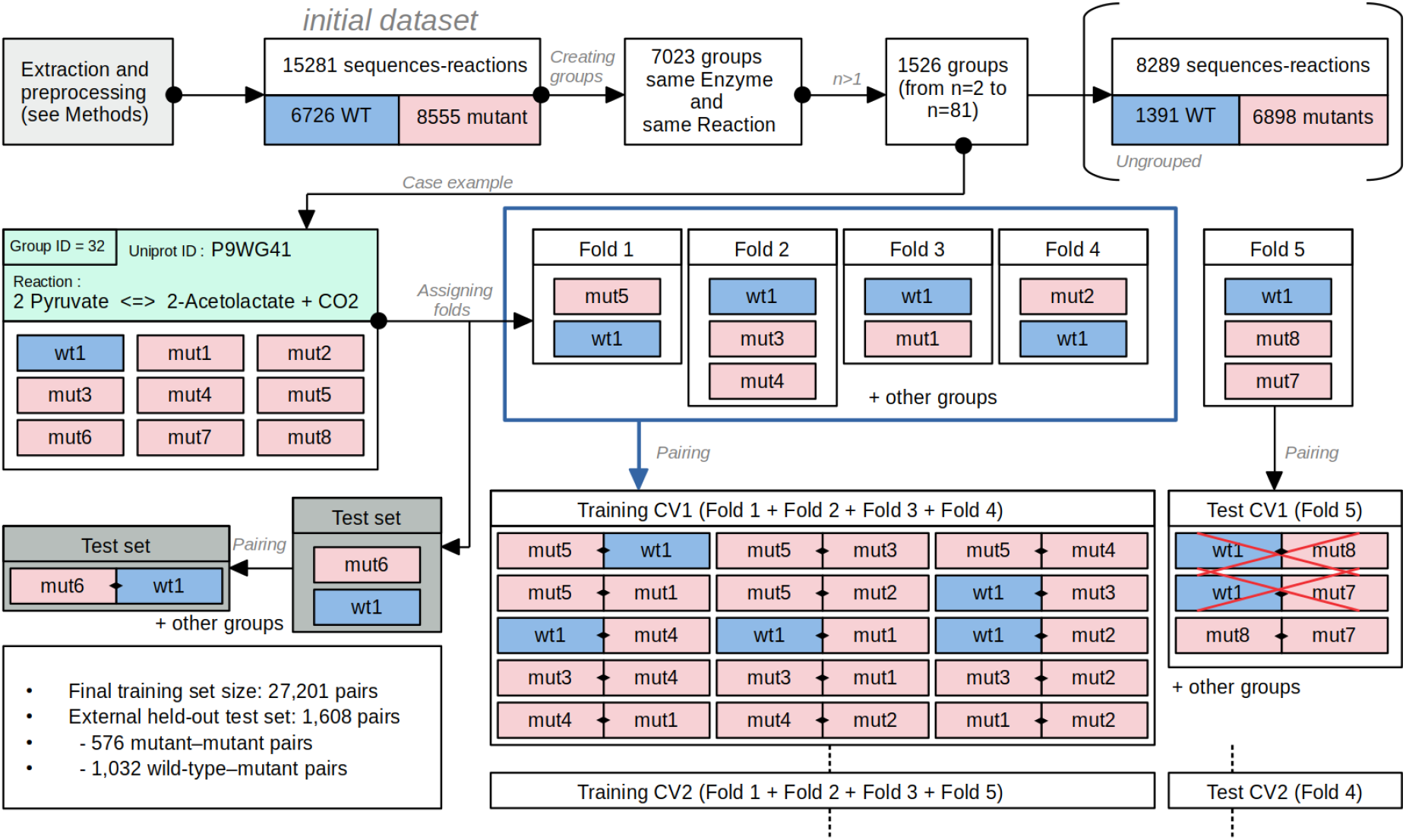
From initial data to cross-validation and external testing. Entries are grouped by enzyme–reaction, and groups containing only a wild-type are removed. Mutants are distributed across five cross-validation folds and one external held-out test split; the wild-type is included in each split containing a mutant from the same group. During cross-validation, four folds are pooled to form all unordered training pairs, *n*(*n* − 1)/2, while the remaining fold provides held-out candidate pairs. For conservative model selection, validation is restricted to mutant–mutant pairs, excluding wild-type–mutant pairs to reduce wild-type anchoring and directional group bias. After hyperparameter selection, the final model is trained on all five CV folds and evaluated once on the external held-out test split. The full external test split contains both mutant–mutant and wild-type–mutant pairs, allowing separate evaluation of the bias-controlled mutant–mutant setting and the less stringent wild-type-anchored setting.

Because pairs are generated within each training split, training sets were substantially larger than a simple sum over singleton entries, and the final training set (after fixing hyperparameters) was larger than the direct sum of CV training splits. On average, each training fold contained ∼ 18,000 pairs and each validation fold ∼ 1,650 pairs. The final aggregated training set comprised 27,201 pairs, while the test set contained 1,608 pairs; when wild-type–mutant pairs were excluded from the test set to avoid bias, this yielded 576 fully unseen mutant–mutant test pairs. Finally, to respect the antisymmetry of pairwise comparisons and to balance the data, we augmented every pair (*a, b*) with its reversed counterpart (*b, a*), using the corresponding ESM-2 representations and the negated target (since Δ_(*b,a*)_ = − Δ_(*a,b*)_). This augmentation enforces the physical constraint that swapping the order of sequences flips the sign of the fold-change, reduces directional bias in the inputs (ESM-2 of sequence *b* and the differential ESM vector), and yields a more balanced, better-conditioned training distribution, which improved generalization in practice.

### Input features and numerical representations

Enzymes were encoded with the ESM-2 protein language model using the publicly available “standard” checkpoint from Hugging Face (33 layers, ∼ 650M parameters)^9^. We also evaluated ESM-1v and ESM-1b as alternative encoders^8,28^; ESM-2 consistently performed best in our setting.

Given a protein sequence of length *T*, the encoder outputs residue-level embeddings **H** ∈ ℝ^*T×d*^, where *d* is the ESM embedding dimension. We compared four pooling operations to obtain a fixed-size sequence representation **v** ∈ ℝ^*d*^. First, mean pooling averages residue embeddings over the sequence,

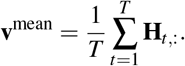

Second, max pooling retains, for each embedding dimension, the largest residue-level value,

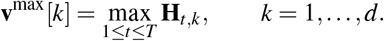

Third, mix pooling provides an intermediate representation between mean and max pooling by averaging the five largest values in each embedding dimension,

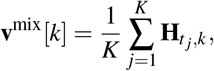

where *K* = 5 and {*t*_1_, …, *t*_5_} are the residue indices corresponding to the 5 largest values of **H**_:,*k*_.

Finally, we introduced absolute max pooling, which retains the most extreme residue-level value in each embedding dimension while preserving its sign:

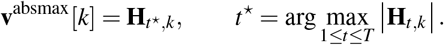

Unlike standard max pooling, this operation also captures large negative activations, which may otherwise be discarded. Absolute max pooling yielded the best downstream performance and was therefore retained as the default pooling operation for FCKcat (Fig. S6).

For an ordered enzyme pair (*a, b*), both amino-acid sequences are encoded independently with ESM-2 and pooled using the same operation. The pairwise enzyme representation is then formed by concatenating the two sequence-level embeddings:

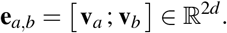

In addition to this concatenated representation, we also evaluated an element-wise difference representation using the bestperforming pooling strategy,

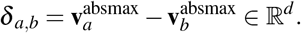

To incorporate reaction information, we append a reaction fingerprint **r** ∈ ℝ^*m*^ derived from the reaction SMILES using RXNFP^29^. RXNFP tokenizes the reaction string and encodes it with a BERT-style transformer trained on USPTO reactions^30^. When reaction information is used together with the enzyme-pair representation, the input vector becomes

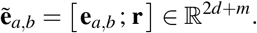

We tested the following feature variants: enzyme-pair concatenation using mean, max, mix, or absolute max pooling; the element-wise difference representation using absolute max pooling; and the concatenation of the absolute max pooling enzyme-pair representation with RXNFP reaction fingerprints to assess whether reaction information improved predictive performance.

### Model training and hyper-parameter optimization

All models were trained using the predefined five-fold splitting strategy and cross-validation described in the Results. To make training symmetric with respect to pair order, each training fold was augmented with the reversed pair (*b, a*) for every pair (*a, b*), with the input order reversed and the sign of the regression target inverted. The regression target was the signed difference in catalytic rates,

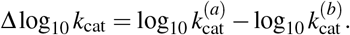

Unless otherwise stated, the default input features corresponded to the concatenated absolute max pooling ESM-2 representations described in the previous subsection,

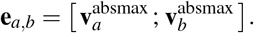

We used gradient-boosted decision trees (XGBoost, GPU backend) with tree_method=gpu_hist and sampling_method=gradient_based. Hyper-parameters were tuned with hyperopt using random search (rand.suggest) over 350 trials on a single NVIDIA RTX 8000 GPU (seed = 42). The search space included learning rate, maximum depth, *𝓁*_2_ and *𝓁*_1_ regularization (reg_lambda, reg_alpha), minimum child weight, *γ*, subsample, column subsample, histogram max_bin, and the number of boosting rounds. Concretely, we sampled

~~~
     *η* ∼ log-uniform(0.01, 0.5), max_depth ∈ {3,…, 12},
     reg_lambda ∼ log-uniform(10^−3^, 10^2^), reg_alpha ∼ log-uniform(10^−3^, 10),
     min_child_weight ∼ log-uniform(0.1, 50), *γ* ∼ log-uniform(10^−4^, 10),
     subsample ∼ uniform(0.5, 1.0), colsample_bytree ∼ uniform(0.5, 1.0),
     max_bin ∈ {128, 192, 256, 320, 384, 448, 512}, num_rounds ∈ [100, 2000].
~~~

For each hyper-parameter setting, we trained on the training partition of each fold and evaluated on the corresponding held-out partition; the objective optimized by hyperopt was the negative mean *R*^2^ across folds, equivalently maximizing mean out-of-fold *R*^2^. The same optimization protocol was applied to all input-feature variants, including concatenated enzyme-pair representations using mean, max, mix, or absolute max pooling; the absolute max pooling element-wise difference representation; alternative ESM encoders; and models with or without RXNFP reaction fingerprints. Unless otherwise noted, the final FCKcat model used concatenated absolute max pooling ESM-2 embeddings.

### *k*_cat_ fold-change notation, reconstruction, and performance metrics

All kinetic values are analyzed on a logarithmic scale. For each measurement, we define *y* := log_10_ *k*_cat_. For any ordered pair of sequences (*a, b*) belonging to the same enzyme–reaction group, we define the log-fold change

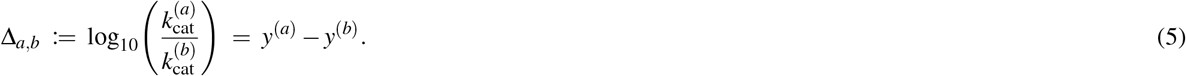

With this convention, swapping the order of sequences flips the sign: Δ_*b,a*_ = − Δ_*a,b*_. When the pair corresponds to a mutant and its wild-type, we use

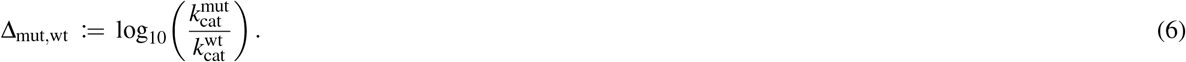

Given a predicted fold change 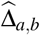 and a known reference measurement, we reconstruct absolute log-values by rearranging Eq. (5):

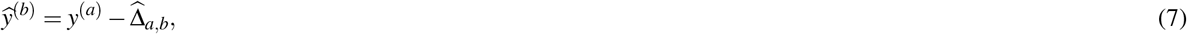

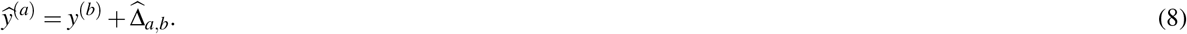

In the wild-type–mutant setting, Eqs. (7)–(8) reduce to reconstructing the unknown log_10_ *k*_cat_ value from the known one using 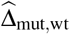, with the sign determined by whether the unknown corresponds to index (*a*) or (*b*).

For *collective prediction*, we use multiple within-group reference measurements, or anchors, to reconstruct the same target and then average. Let *j* denote a target sequence, typically a test mutant, in enzyme–reaction group *g*, and let *𝒜*_*g*_ be the set of available anchor sequences in the same group with experimental measurements 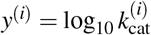. In our setting, *𝒜*_*g*_ corresponds to the training sequences from group *g*. For each anchor *i* ∈ *𝒜*_*g*_, we predict the pairwise fold change 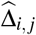 and reconstruct an anchor-specific estimate of *y*^(*j*)^ via

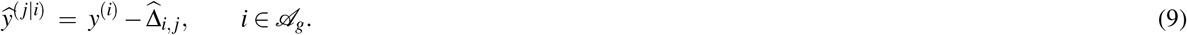

We then define the collective estimate as the average across anchors,

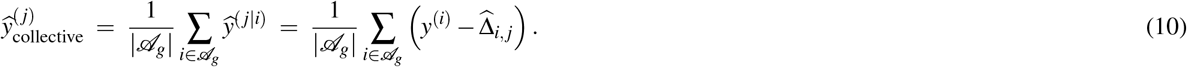

When desired, 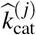 is obtained by inverse transformation: 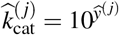. In practice, collective prediction is only applied when |*𝒜*_*g*_| *>* 0.

To quantify how much apparent performance can be explained by enzyme–reaction group-level shortcuts, we implemented two baselines. First, for absolute log_10_ *k*_cat_, we predict for each test entry *e* in group *g* the mean of the training values from the same group:

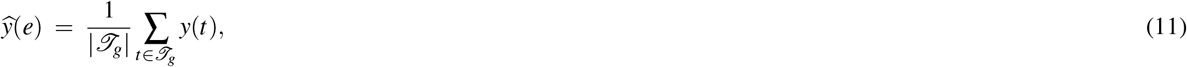

where *𝒯*_*g*_ denotes all training entries, wild-type and mutant, in group *g*. Second, for mutant–wild-type fold changes, we compute for each group *g* the mean training effect

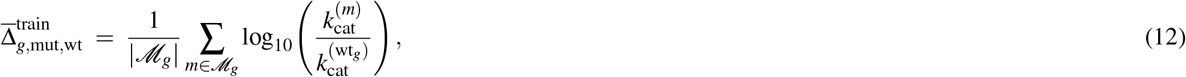

where ℳ_*g*_ is the set of training mutants in group *g*, and wt_*g*_ denotes the corresponding wild-type. We then predict for a test mutant *m* ∈ *g*

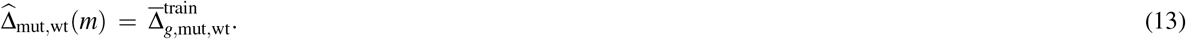

Groups without any training mutants, i.e. |*M*_*g*_| = 0, were excluded from the group-average Δ_mut,wt_ baseline and from analyses that depend on 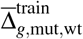.

For regression, we report the coefficient of determination *R*^2^ and, where indicated, Pearson’s correlation coefficient *r* between predicted and experimentally observed values. Given true targets *z*_*i*_ and predictions 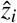 over *n* evaluated examples, we compute

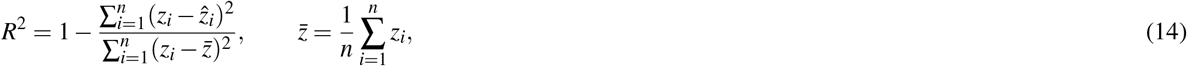

This is a predictive *R*^2^ computed from the raw model predictions 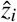, i.e. relative to the ideal identity relationship 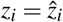, and not the coefficient of determination of a post hoc linear fit between *z*_*i*_ and 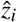.

We further compute Pearson’s correlation coefficient as

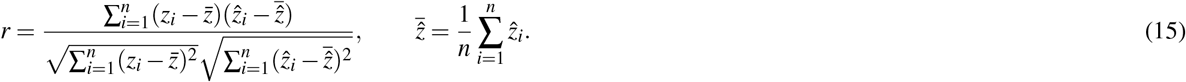

Unless stated otherwise, *R*^2^ is computed on the pairwise fold-change target Δ_*a,b*_, defined in Eq. (5).

To assess the ability to predict the sign of mutation effects on sequence pairs, we report sign accuracy and Matthews correlation coefficient (MCC). With TP, TN, FP, and FN denoting the usual confusion-matrix counts, we compute

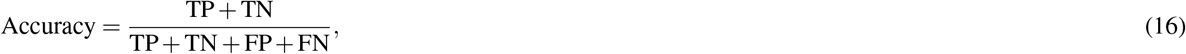

and

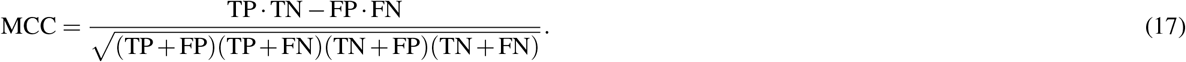

We additionally evaluate these quantities as a function of predicted effect size by applying a magnitude threshold δ and retaining only pairs with 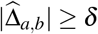.

To isolate mutation-level signal beyond group-level averages and to avoid inflated pooled performance dominated by between-group differences, we perform a within-group residual analysis. For each group *g* with |*M*_*g*_| *>* 0, we compute 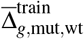 using Eq. (12) and define residuals for each test mutant–wild-type pair as

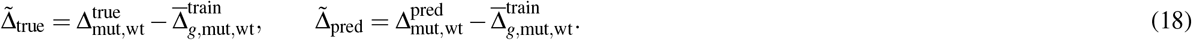

Under this transformation, the group-average baseline collapses to 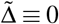 by construction, so any correlation between 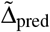 and 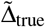 reflects mutation-specific information beyond group effects.

### Evaluation protocol

Models were trained and selected using the cross-validation (CV) splits. Our primary goal is to measure mutation-level generalization while minimizing shortcuts induced by enzyme–reaction group structure. We therefore distinguish three evaluation settings: (i) *unseen mutant–mutant* pairs, where both sequences in each evaluated pair are mutants and neither sequence appears in the training folds; (ii) *wild-type–mutant* pairs, where a mutant is compared to its wild-type reference; and (iii) the *full test set*, which includes both pair types. Hyper-parameters were selected by maximizing mean out-of-fold *R*^2^ on the held-out CV fold, and the final model was refit on the pooled training folds and evaluated once on an external test split.

To reduce wild-type anchoring and directional group biases during model selection, our primary (most conservative) protocol evaluates only mutant–mutant pairs in validation and test. Concretely, after mutants were distributed across CV folds and the external test set within each enzyme–reaction group, we formed all unordered sequence pairs *within each split* to create training/validation/test pairs. For the conservative evaluation, all wild-type–mutant pairs were removed from the validation and test splits, so that every evaluated pair involves two mutants that are both unseen during training.

Although wild-type–mutant pairs are excluded from validation/test in the conservative protocol, wild-type sequences are retained within the splits to (i) preserve the natural pairing structure of mutant datasets and (ii) enable downstream analyses that reconstruct absolute *k*_cat_ from fold changes using experimental anchors. We therefore also report performance on wild-type– mutant pairs and on the full test set, but interpret these settings as less stringent because the wild-type provides group-level information that can inflate apparent performance.

### Leakage-controlled benchmarking against existing models

To compare against existing predictors (DLKcat and EITLEM), we constructed leakage-free, task-matched test intersections. For each external model, we obtained the original training and test splits released by the authors and defined training exposure as any amino-acid sequence appearing in the corresponding training data. We then removed from the external test split any entries whose sequence occurs in any of the external model’s training sets; for EITLEM, we additionally treated sequences appearing in auxiliary training tasks used for transfer learning (e.g., *K*_*M*_ and *k*_cat_*/K*_*M*_) as training exposure. Under this definition, approximately 77% of sequences in the published EITLEM *k*_cat_ test set appear in its *k*_cat_ training set, increasing to ∼ 90% when including auxiliary-task training data; additionally, ∼ 25% of the *k*_cat_ test entries match a training entry with the same sequence and identical *k*_cat_ value and differ only in substrate annotation. After leakage removal, we formed the intersection between the corrected external test set and our own held-out test set, and evaluated all methods on exactly the same entries. Because each external model provides different published test sets and the leakage-filtering step removes a different fraction of entries, the resulting intersection sizes (and thus the number of evaluated pairs) differ between the FCKcat-DLKcat and FCKcat-EITLEM comparisons. For fold-change comparisons, we restricted evaluation to mutant–mutant pairs and computed fold changes from predicted absolute values when necessary, using the same definition of Δ_*ab*_ as in Eq. (5). Performance on these intersections is reported using Pearson’s correlation coefficient *r* for regression and sign accuracy and Matthews correlation coefficient (MCC) for direction-of-change classification.

## Code availability

The Python code used to generate all results is publicly available at https://github.com/yvanrousset/FCKcat

## Acknowledgements

Computational infrastructure and support were provided by the Centre for Information and Media Technology at Heinrich Heine University Düsseldorf.

## Funding Statements

This work was funded through grants to M.J.L. by the European Union (ERC AdG “MechSys”–Project ID 101055141) and by the Deutsche Forschungsgemeinschaft (DFG, German Research Foundation: CRC 1310, and, under Germany’s Excellence Strategy, EXC 2048/2–Project ID 390686111).

## Competing interests

A.K. and M.J.L. are founders and board members of ProVis ProteinVision GmbH.

## Notes

### Competing Interest Statement

The authors have declared no competing interest.

### Summary of Updates

Code availability Acknowledgements Conflict of interest Figures Figures' legend

