## Supplementary Information for "Overcoming systematic data biases enables accurate prediction of enzyme *k*_cat_ fold-changes for computational protein design"

### Supplementary Figures

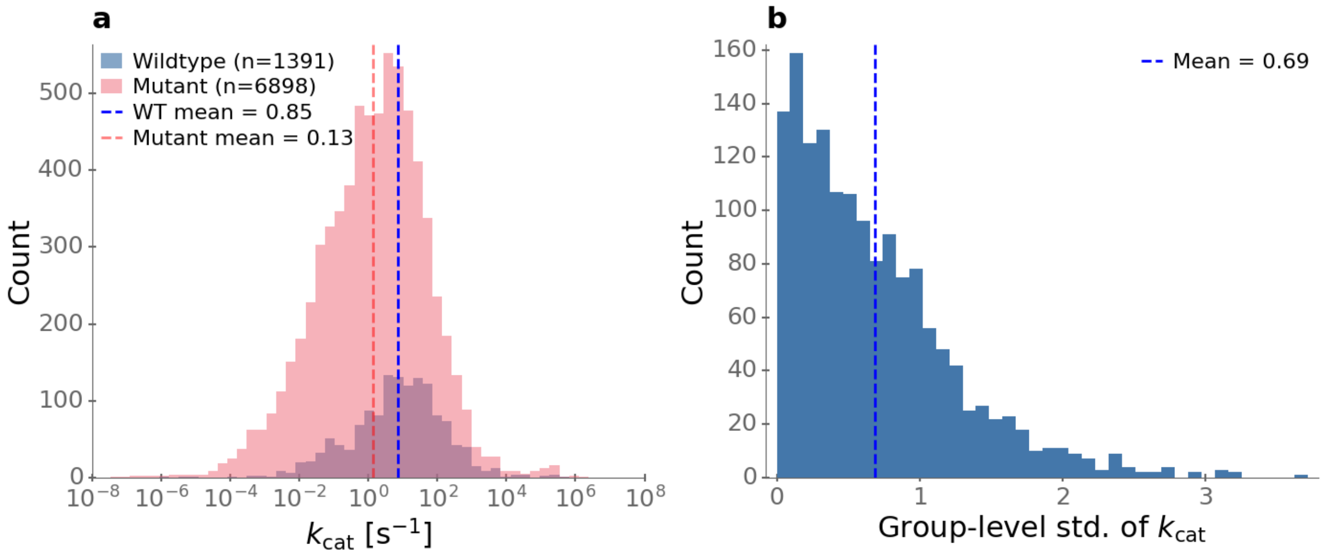


**Figure S1. Within-group variation in $\log_{10} k_{cat}$ is much lower than between-group variation. a,** Distributions of wild-type and mutant $k_{cat}$ values. Mean $\log_{10} k_{cat}$ is 0.85 for wild-types and 0.13 for mutants. **b,** Distribution of within-group standard deviations of $\log_{10} k_{cat}$; the mean within-group standard deviation is 0.69.

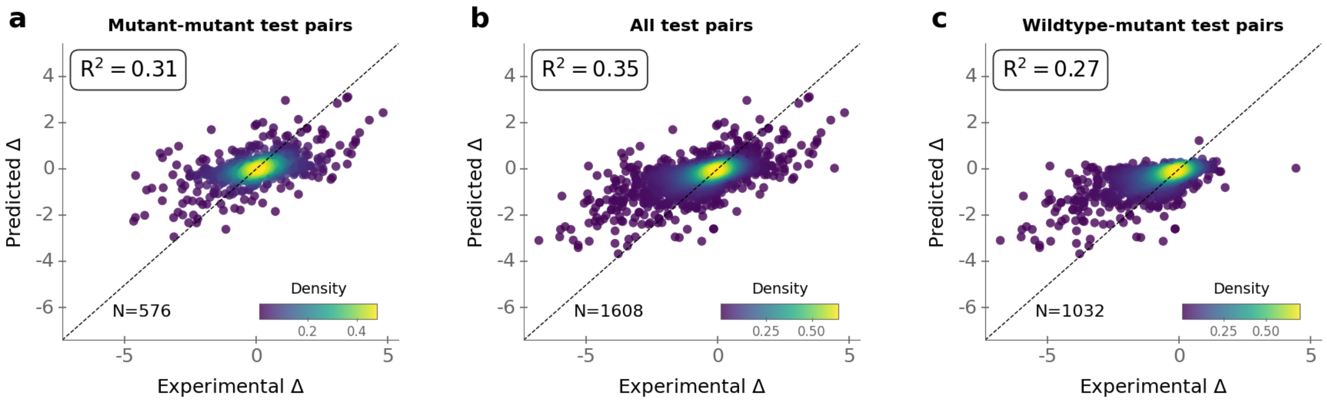


**Figure S2. Apparent performance increases with the inclusion of wild-type pairs due to reintroduction of group-level bias. a,** Evaluation on mutant–mutant pairs only, corresponding to the bias-controlled setting. **b,** Evaluation on all sequence pairs. **c,** Evaluation restricted to mutant–wild-type pairs.

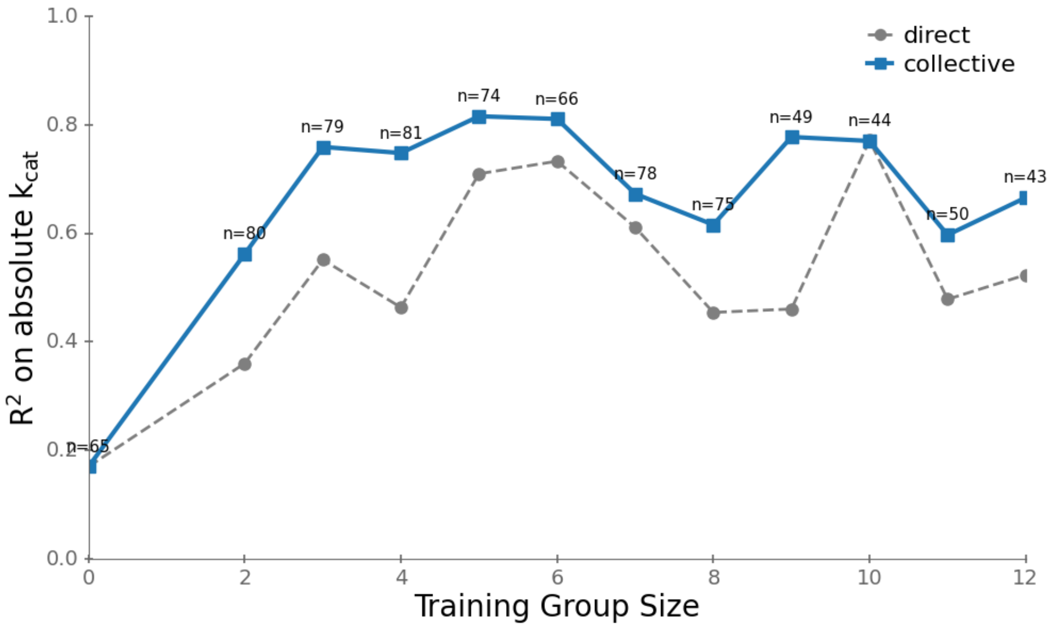


**Figure S3. Prediction accuracy of collective predictions increases with training group size.** Prediction accuracy ($R^2$ on $\log_{10} k_{\mathrm{cat}}$) for unseen mutants as a function of the number of mutant reference sequences available in the training set for each enzyme–reaction group.

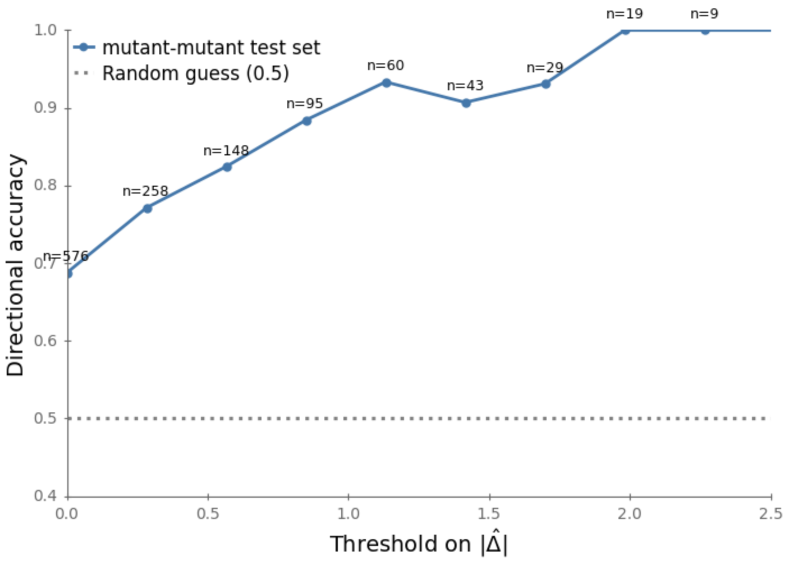


**Figure S4. Directional predictions become highly reliable for large predicted effect sizes.** For a threshold $\delta$ on the predicted magnitude of the log-fold change, we evaluate only variants with $|\widehat{\Delta}_{\text{mut,mut}}| \geq \delta$. The sign accuracy increases from 69% at $\delta = 0$ to 90% for $\delta > 1.0$, corresponding to predicted $k_{\text{cat}}$ changes exceeding one order of magnitude.

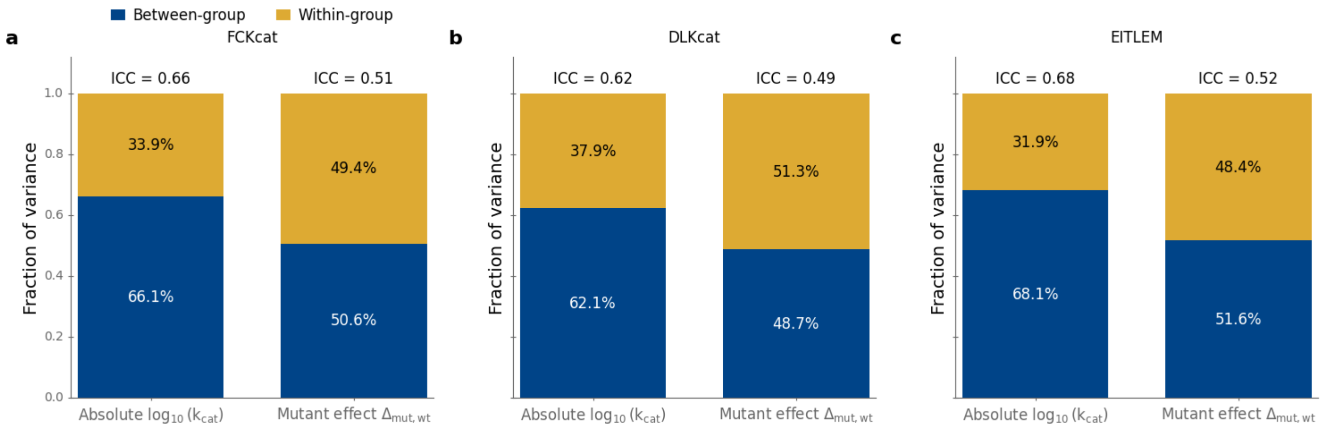


**Figure S5.** **The partitioning between between-group and within-group variance is consistent across $k_{cat}$ datasets.** Stacked bars show the fraction of variance attributable to between-group differences and within-group variation, estimated with random-intercept linear mixed-effects models using enzyme group identity. Group identity explains a substantial fraction of the variance in log-transformed $k_{cat}$ values as well as in mutant effects relative to wild-type across the datasets used in this study, DLKcat, and EITLEM. As the EITLEM and DLKcat datasets do not provide explicit enzyme–reaction identifiers, we approximated their enzyme groups by clustering protein sequences using a RapidFuzz-based procedure[1] with a 98% sequence-similarity threshold.

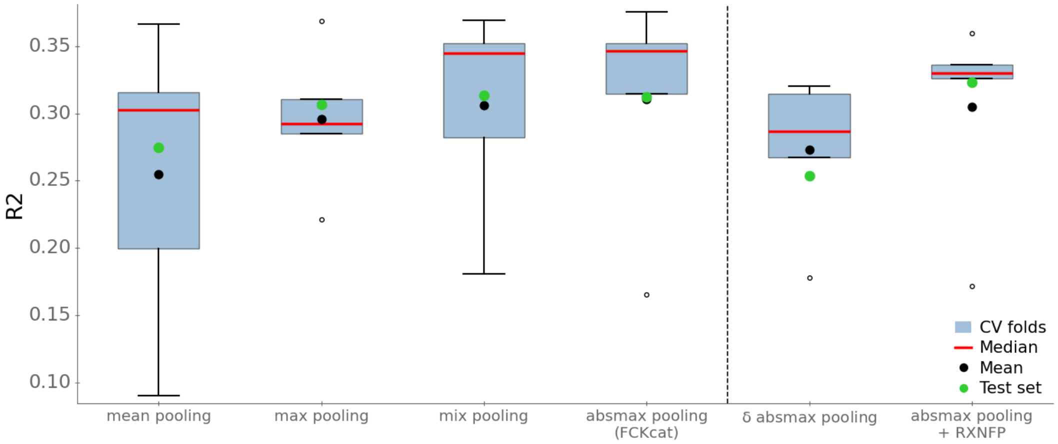


**Figure S6. Feature-set comparison by cross-validation.** Per-fold performance of different input representations, evaluated using the coefficient of determination ($R^2$). The red horizontal line indicates the median across folds, the black dot indicates the mean, and the green dots denote performance on the held-out test set.
